# Structures of α-synuclein filaments from multiple system atrophy

**DOI:** 10.1101/2020.02.05.935619

**Authors:** Manuel Schweighauser, Yang Shi, Airi Tarutani, Fuyuki Kametani, Alexey G. Murzin, Bernardino Ghetti, Tomoyasu Matsubara, Taisuke Tomita, Takashi Ando, Kazuko Hasegawa, Shigeo Murayama, Mari Yoshida, Masato Hasegawa, Sjors H.W. Scheres, Michel Goedert

## Abstract

Synucleinopathies are human neurodegenerative diseases that include multiple system atrophy (MSA), Parkinson’s disease, Parkinson’s disease dementia (PDD) and dementia with Lewy bodies (DLB) (1). Existing treatments are at best symptomatic. These diseases are characterised by the presence in brain cells of filamentous inclusions of α-synuclein, the formation of which is believed to cause disease (2, 3). However, the structures of α-synuclein filaments from human brain are not known. Here we show, using electron cryo-microscopy, that α-synuclein inclusions from MSA are made of two types of filaments, each of which consists of two different protofilaments. Non-proteinaceous molecules are present at the protofilament interfaces. By two-dimensional class averaging, we show that α-synuclein filaments from the brains of patients with MSA and DLB are different, suggesting that distinct conformers (or strains) characterise synucleinopathies. As was the case of tau assemblies (4–9), the structures of α-synuclein filaments extracted from the brains of individuals with MSA differ from those formed *in vitro* using recombinant proteins, with implications for understanding the mechanisms of aggregate propagation and neurodegeneration in human brain. These findings have diagnostic and potential therapeutic relevance, especially in view of the unmet clinical need to be able to image filamentous α-synuclein inclusions in human brain.

---

A causal link between α-synuclein assembly and disease was established by the findings that missense mutations in *SNCA*, the α-synuclein gene, and multiplications thereof, give rise to rare inherited forms of Parkinson’s disease and PDD (3, 10). Some mutations also cause DLB. Missense mutations G51D (11, 12) and A53E (13) in *SNCA* can give rise to atypical synucleinopathies, with mixed Parkinson’s disease and MSA pathologies. Sequence variation in the regulatory region of *SNCA* is associated with increased expression of α-synuclein and heightened risk of developing idiopathic Parkinson’s disease (14), which accounts for over 90% of disease cases. In addition, overexpression of human mutant α-synuclein in animal models causes its assembly and neurodegeneration (15).

MSA is a sporadic synucleinopathy of adult onset (16). Patients suffer from parkinsonism, cerebellar ataxia and autonomic failure (17–20). Cases of MSA are classified as MSA-P, with predominant parkinsonism caused by striatonigral degeneration, and MSA-C, with cerebellar ataxia associated with olivopontocerebellar atrophy.

Autonomic dysfunction (Shy-Drager syndrome), which includes orthostatic hypotension and urogenital symptoms, is common to MSA-P and MSA-C. Neuropathologically, MSA is defined by regional nerve cell loss and the presence of abundant filamentous α-synuclein inclusions in oligodendrocytes: glial cytoplasmic inclusions (GCIs) or Papp-Lantos bodies (21–25). Smaller numbers of α-synuclein inclusions are also present in nerve cells (26, 27). The formation of α-synuclein assemblies is an early event that precedes neurodegeneration (28–30). Mean disease duration is 6-10 years, but survival times of 18-20 years have been reported (31). Late appearance of autonomic dysfunction correlates with prolonged survival (32).

α-Synuclein is a 140 amino acid protein, over half of which consists of seven imperfect repeats, with the consensus sequence KTKEGV (residues 7-87). They encompass the lipid-binding domain (33). The repeats partially overlap with a hydrophobic region of α-synuclein (residues 61-95), also known as the non-β-amyloid component (NAC) (34). The NAC is necessary for assembly of recombinant α−synuclein into filaments (35–37). The carboxy-terminal region of α-synuclein (residues 96-140) is negatively charged and truncation of this region has been linked to increased assembly into filaments (38). Upon assembly, recombinantly expressed α-synuclein undergoes conformational changes and takes on a cross-β structure characteristic of amyloid (39, 40). Using a variety of techniques, the cores of α-synuclein filaments extracted from cerebellum of MSA patients or assembled from recombinant protein *in vitro* have been shown to encompass 70 amino acids, extending approximately from residues 30-100 (41–44).

Seeded assembly of α-synuclein, propagation of inclusions and nerve cell death have been demonstrated in a variety of systems (45–48). Assemblies of recombinant α-synuclein with different morphologies displayed distinct seeding capacities (49). Moreover, GCI α-synuclein has been reported to be approximately three orders of magnitude more potent than Lewy body α-synuclein in seeding aggregation of α-synuclein (50). Indirect evidence has also suggested that distinct conformers of assembled α-synuclein may characterise MSA and disorders with Lewy pathology (51–56). Solubility in SDS distinguishes α-synuclein filaments of MSA from those of DLB (57).

Progress in understanding synucleinopathies and the existence of molecular conformers is hampered by the lack of atomic structures of α-synuclein filaments from human brain. Here we used electron cryo-microscopy (cryo-EM) to determine high-resolution structures of the cores of α-synuclein filaments that were extracted from the brains of individuals with MSA. In addition, we used two-dimensional (2D) class averaging to show that α-synuclein filaments from MSA and DLB are different.

## NEUROPATHOLOGICAL CHARACTERISTICS

α-Synuclein filaments were extracted using sarkosyl from the putamen of 5 individuals with a neuropathologically confirmed diagnosis of MSA. For MSA cases 1 and 3, filaments were also extracted from cerebellum and frontal cortex. Most sarkosyl-insoluble α-synuclein phosphorylated at S129 was soluble in SDS. More than 90% of filamentous α-synuclein inclusions are phosphorylated at S129 (58). One case (number 1) was diagnosed as MSA-P (age at death 85 years) and four cases (numbers 2-5) as MSA-C (ages at death 68, 59, 64 and 70 years). Disease durations for MSA cases 1-5 were: 9, 18, 9, 10 and 19 years.

Abundant GCIs and neuronal inclusions were stained by an antibody specific for α-synuclein phosphorylated at S129 [Figure 1a; Extended Data Figure 1a,b]. By negative-stain electron microscopy (EM), all 5 cases of MSA showed a majority of twisted filaments, with a diameter of 10 nm and a periodicity of 80-100 nm [Figure 1b; Extended Data Figure 1c,d]. Immunogold negative-stain EM with anti-α-synuclein antibody PER4 showed decoration of MSA filaments [Extended Data Figure 2], in confirmation of previous findings using different cases of MSA (23). By immunoblotting of sarkosyl-insoluble material with Syn303 and PER4, full-length α-synuclein, as well as high-molecular weight aggregates, were in evidence. High-molecular weight aggregates were present in larger amounts relative to full-length, monomeric protein, when Syn303 was used for immunoblotting. Moreover, truncated α-synuclein was also present, especially with Syn303. When antibody pS129 was used, full-length α-synuclein was the predominant species [Extended Data Figure 1e]. Consistent with immunostaining [Figure 1a], sarkosyl-insoluble fractions from the putamen of MSA cases 1 and 3 contained lower levels of α-synuclein than those from MSA cases 2, 4 and 5.

**Figure 1.**
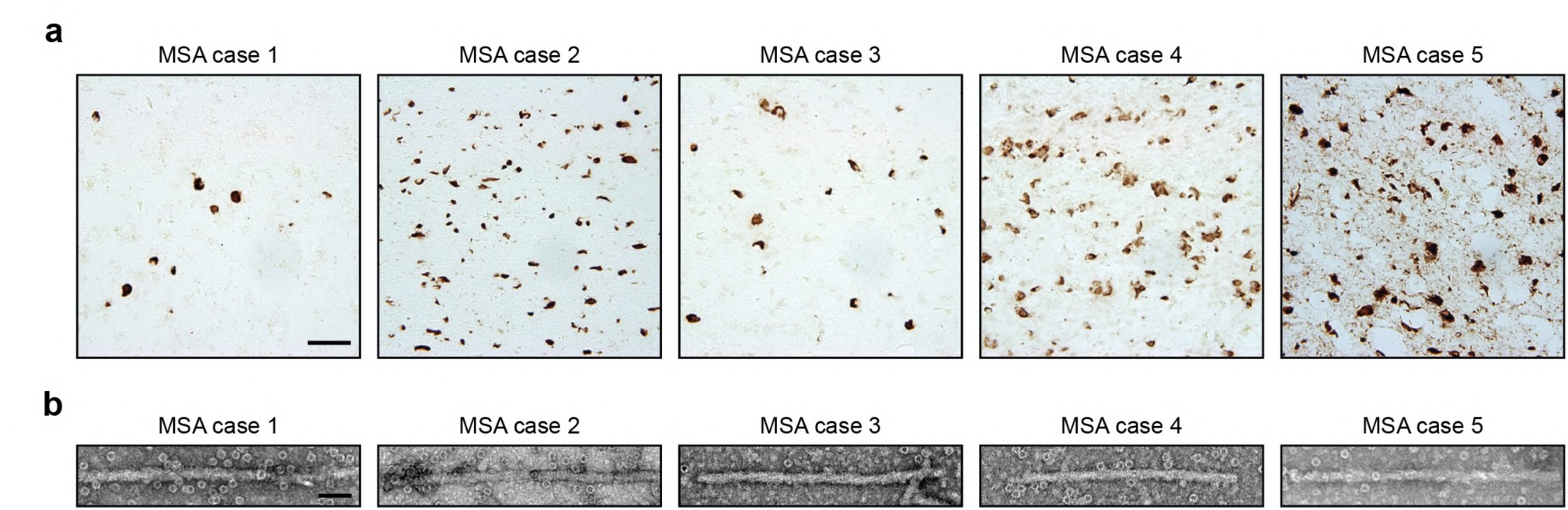
Filamentous α-synuclein pathology of MSA. (a), Staining of neuronal and glial inclusions in the putamen of MSA cases 1-5 by an antibody specific for α-synuclein phosphorylated at S129 (brown). Scale bar, 50 μm. (b), Negative stain electron micrographs of filaments from the putamen of MSA cases 1-5. Scale bar, 50 nm.

Seeded aggregation of expressed wild-type human α-synuclein was observed in SH-SY5Y cells after addition of sarkosyl-insoluble seeds from the putamen of MSA cases 1-5 [Extended Data Figure 3]. Seeds from MSA case 3 were the most potent, whereas those from MSA case 2 were least effective at inducing seeded aggregation. Seeds from MSA cases 1, 4 and 5 gave similar results.

**Figure 2.**
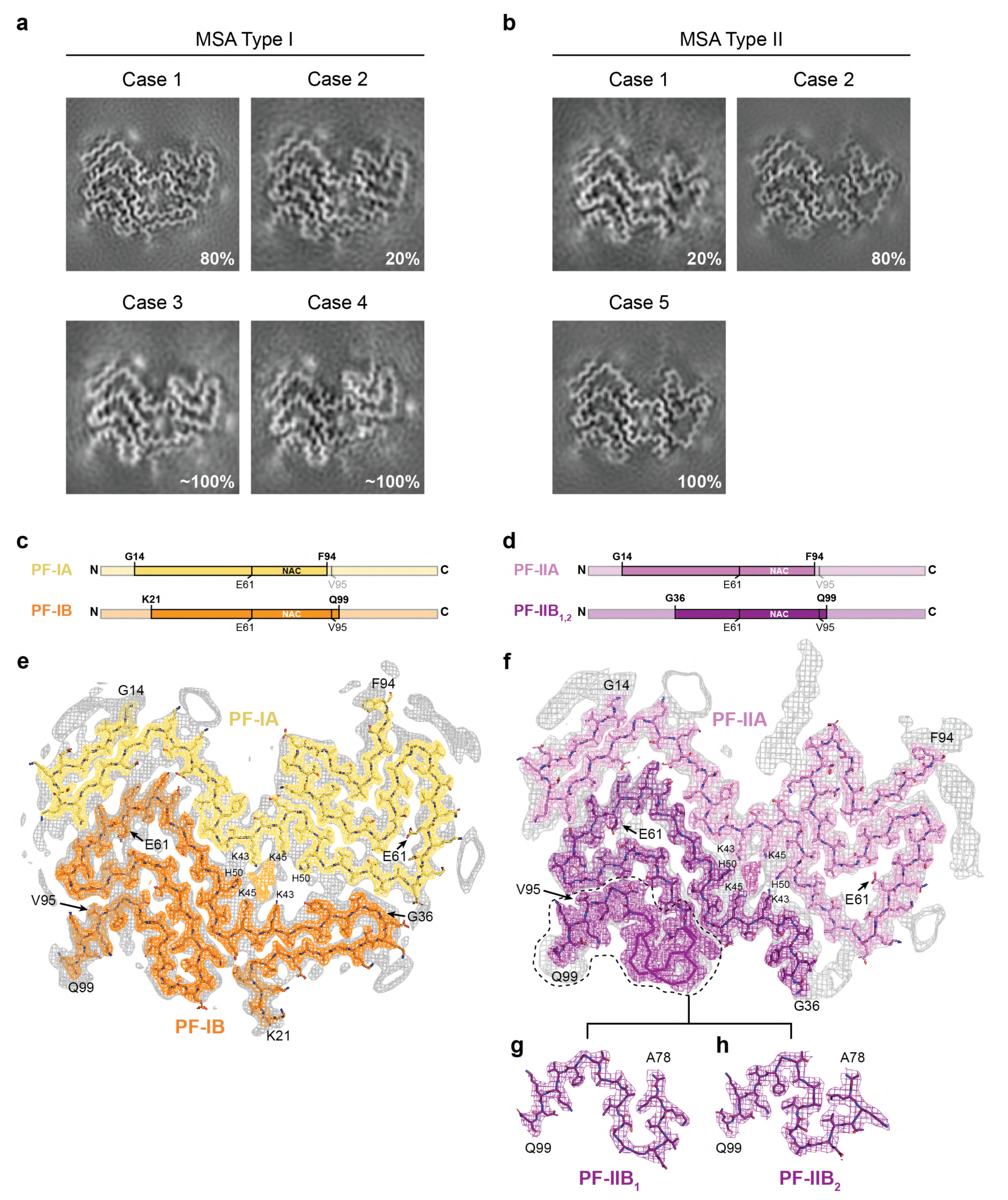
Cryo-EM maps and atomic models of MSA Type I and Type II α-synuclein filaments. (a,b), Cryo-EM maps of Type I filaments from the putamen of MSA cases 1, 2, 3 and 4 (a) and of Type II filaments from the putamen of MSA cases 1, 2 and 5 (b). (c,d), Schematic of the primary structure of human α-synuclein, indicating the cores of protofilaments (PF) IA, IB, IIA and IIB. The NAC domain (residues 61-95) is also shown. (e,f), Sharpened, high-resolution cryo-EM maps of MSA Type I (e) and Type II (f) α-synuclein filaments with overlaid atomic models. Unsharpened, 4.5 Å low pass-filtered maps are shown in grey. They show weaker densities that extend from the N- and C-terminal regions, a peptide-like density in PF-IIA, as well as weaker densities bordering the solvent-exposed chains of K32 and K34 in PF-IA, PF-IB and PF-IIA. Weaker densities bordering the solvent-exposed chains of K58 and K60 in PF-IA and PF-IIA are also present. (g,h), Cryo-EM structures of A78-Q99 of PF-IIB, illustrating heterogeneity (PF-IIB_1_ and PF-IIB_2_). Please note the strong density at the protofilament interfaces of MSA Type I and Type II filaments. It is surrounded by the side chains of K43, K45 and H50 from each protofilament.

**Figure 3.**
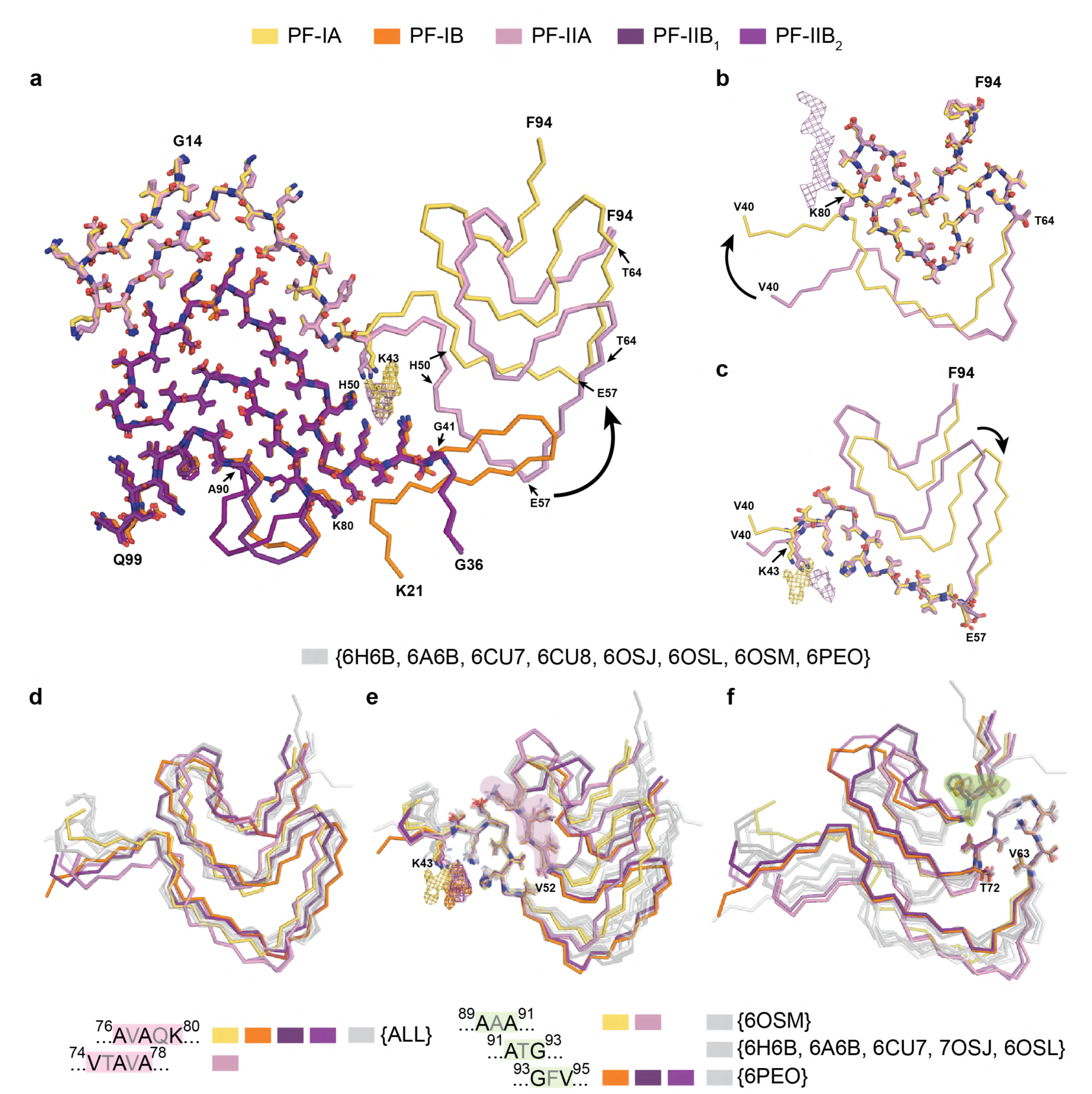
Comparison of MSA α-synuclein protofilament folds. (a), Overlay of the structures of MSA PF-IA, PF-IB, PF-IIA and PF-IIB. The black arrow indicates the direction of the conformational change that occurs at K43 of PF-IA and PF-IIA.(b,c), Three-layered L-shaped motifs of PF-IA (yellow) and PF-IIA (pink) are aligned, based on the structural similarities between T64-F94 (b) and T44-E57 (c). Black arrows indicate the direction of the conformational change that occurs at T64 (b) or E57 (c) of PF-IA and PF-IIA. The peptide-like density in PF-IIB is shown as a pink mesh. (d), Overlay of the three-layered L-shaped motifs of MSA α-synuclein filaments and filaments assembled *in vitro* using recombinant α-synuclein. (e), Overlay of MSA and recombinant α-synuclein structures based on the conserved turn at K43-V52, revealing a conserved interface between E46-V49 and V74-A78 or A76-K80 (red highlight). (f), Overlay of MSA and recombinant α-synuclein structures based on the conserved turn at V63-T72, revealing a conserved interface between A69-T72 and A89-91, A91-G93 or G93-V95 (green highlight).

## CRYO-EM REVEALS TWO TYPES OF α-SYNUCLEIN FILAMENTS IN MSA

Sarkosyl-insoluble filaments from all 5 cases of MSA were imaged by cryo-EM [Extended Data Figure 4a-c]. They looked identical upon visual inspection of the micrographs, but the existence of two types (Type I and Type II) of filaments was revealed by reference-free 2D class averages [Extended Data Figure 4d-f]. Type I filaments were less symmetrical than Type II filaments. In putamen, Type I:II filament ratios were 80:20 for MSA case 1 (disease duration of 9 years) and 20:80 for MSA case 2 (disease duration of 18 years). MSA cases 3 and 4 (disease durations of 9 and 10 years) had mostly Type I filaments, whereas MSA case 5 (disease duration of 19 years) had only Type II filaments (Figure 2a,b). This suggests that the duration of MSA may correlate with the ratio of filament types in putamen, but additional cases of disease are required to establish this more firmly. However, what is the case in putamen may not be true of α-synuclein filaments from other affected brain regions. Whereas we identified predominantly Type I filaments in the putamen of MSA cases 1 and 3 [Figure 2], we found almost exclusively Type II filaments in the cerebellum of MSA case 1 and in the frontal cortex of MSA case 3 [Extended Data Figure 4a]. It remains to be seen if MSA Type I and Type II α-synuclein filaments are common to both nerve cells and glial cells.

## MSA TYPE I AND TYPE II PROTOFILAMENTS ADOPT EXTENDED FOLDS

We used helical reconstruction in RELION to determine the structures of MSA Type I and Type II filaments from putamen to resolutions sufficient for *de novo* atomic modelling [Figure 2, Extended Data Table 1]. The best structures were resolved to resolutions of 2.6 Å for Type I filaments from MSA case 1 and 3.1 Å for Type II filaments from MSA case 2 [Extended Data Figure 5]. MSA Type I and Type II filaments are each made of two protofilaments, which consist of an extended N-terminal arm and a compact C-terminal body [Figure 2; Extended Data Figure 6]. Type I and Type II filaments are asymmetrical, with one protofilament being larger than the other. PF-IA, the larger protofilament of MSA Type I filaments, comprises residues G14-F94 of α-synuclein, whereas PF-IB, the smaller protofilament, consists of residues K21-Q99 (Figure 2c). In MSA Type II filaments, PF-IIA and PF-IIB comprise residues G14-F94 and G36-Q99, respectively (Figure 2d). Protofilament folds differ from each other within and between filament types. MSA Type I and Type II filaments are thus made of four distinct protofilaments [Figure 2; Figure 3a].

The N-terminal arm of PF-IA consists of a cross-β hairpin spanning G14-G31 and an extended one-layered L-shaped motif at K32-K45. The C-terminal body of PF-IA adopts a three-layered L-shaped motif. The outer layer is the longest, comprising E46-V66. It packs against the outer side of the central layer, which comprises G67-E83. A salt bridge between E46 and K80 stabilises the interaction between these two layers. The shorter inner layer comprises G84-F94 and packs against the other side of the central layer. Pairs of glycines are present in both turns that connect the layers. PF-IA comprises 12 β-strands.

The N-terminal arm of PF-IB consists only of a cross-β hairpin at G25-K45. The three-layered L-shaped motif of its C-terminal body is topologically similar to that in PF-1A. Nevertheless, the two motifs differ in structure, most notably in the packing of the inner layer against the central layer by the residues following G86. Whereas the body of PF-IA ends at F94, that of PF-IB extends to Q99. PF-IB comprises 10 β-strands.

Like PF-IA, PF-IIA is made of residues G14-F94 of α-synuclein and both protofilaments have a similar N-terminal arm [Figure 3b,c]. Although the C-terminal body also adopts a three-layered L-shaped motif, its conformation differs in PF-IIA. Whereas G47-V52 from the outer layer pack against A76-K80 from the central layer in Type I protofilaments, in PF-IIA this packing is shifted by two residues and involves V74-A78 in the central layer. This change in packing creates a sizeable cavity between the central layer and the L-shaped bend at E57 in the outer layer. It increases the distance between the Cα atoms of E46 and K80 by 5 Å, but a salt bridge may still form between their side chains. Like PF-IA, PF-IIA also comprises 12 β-strands.

PF-IIB, which extends from G36-Q99, is the smallest protofilament core and comprises 9 β-strands. Its N-terminal arm is made of a single L-shaped conformation at G36-K45. Its C-terminal body forms a three-layered L-shaped motif, which exists in two conformations: PF-IIB_1_, which is virtually identical to PF-IB, and PF-IIB_2_, which has a different backbone conformation at T81-A90 [Figure 2f, inset; Extended Data Figure 7]. Based on the number of classified helical segments, the ratio of Type II_1_ : Type II_2_ filaments was 20:80 [Extended Data Table 1].

## MSA PROTOFILAMENTS PACK ASYMMETRICALLY AND ENCLOSE ADDITIONAL MOLECULES

In MSA Type I and Type II α-synuclein filaments, two non-identical protofilaments pack against each other through an extended interface [Figure 2; Extended Data Figure 6]. In both types, the packing of Q24-G41 from the L-shaped N-terminal arm of PF-A against PF-B effectively adds a fourth layer to the L-shaped motif of PF-B. The interfaces of both filaments form a large cavity that is surrounded in a pseudo-symmetrical manner by the side chains of K43, K45 and H50 from each protofilament. This cavity encloses an additional strong density that is not connected to the protein density [Figure 2; Extended Data Figure 5]. In Type I filaments, this cavity is larger than in Type II filaments and contains additional, smaller densities between H50, G51 and A53 of PF-IA, and between V37 and Y39 of PF-IB. On this side of the cavity, where the N-terminal arm of PF-B packs against the outer layer of the L-shaped motif of PF-A, the protofilament interfaces of MSA Type I and Type II filaments are most different.

Because of variations in the height of both polypeptide chains along the helical axis, each α-synuclein molecule interacts with three different molecules in the opposing protofilament. If one considers the interaction between two opposing molecules to be on the same rung in the central cavity, the N-terminal arm of PF-A interacts with the C-terminal body of the PF-B molecule, which is one β−sheet rung higher, while the C-terminal body of PF-IA interacts with the N-terminal arm of the PF-IB molecule, which is one rung lower [Extended Data Figure 8].

The chemical nature of the density, which is surrounded by K43, K45 and H50 from each protofilament, remains to be established. The observations that it is disconnected from the density of the α-synuclein polypeptide chains, and that it would need to compensate 4 positive charges for every β-sheet rung, suggest that this density is non-proteinaceous.

Besides the density in the large cavity at the protofilament interface, several other densities are visible at lower intensities. At the N- and C-termini of the ordered cores of all four protofilaments, fuzzy densities probably correspond to less well-ordered extensions of the core. The longest extensions are seen for PF-IA and PF-IIA. Unlike PF-IA, a peptide-like density of unknown identity is packed against residues K80-E83 of PF-IIA. It may correspond to an extension of α-synuclein at the C-terminus of PF-IIA, or to an unknown protein that is bound to the filament core. Additional unconnected densities are observed in front of pairs of lysines on the exterior of the filaments, i.e. in front of K32 and K34 of PF-IA, PF-IB and PF-IIA, as well as in front of K58 and K60 of PF-IA and PF-IIA. Similar densities were observed in front of pairs of lysines on the exterior of tau filaments from Alzheimer’s disease (4, 6), Pick’s disease (5), CTE (7) and CBD (9). The identities of the molecules that form these densities remains unknown. To compensate for the positive charges of lysines, these densities must be negatively charged. A smaller additional density, which may correspond to a positively charged molecule or an ion, is visible between the closely packed, negatively charged side chains of E57 from PF-IA and E35 from PF-IB.

In the structures of MSA Type I and Type II α-synuclein filaments, residues G51 and A53 reside in the protofilament interfaces, close to K43, K45 and H50. Mutations G51D and A53E in *SNCA*, which cause familial Parkinson’s disease, can give rise to mixed MSA and Parkinson’s disease pathologies (11–13). They are the only known disease-causing mutations that increase the negative charge of α-synuclein. In PF-IA and PF-IB, there is room for the side chain of D or E. Similarly, in PF-IIA and PF-IIB, small cavities can accommodate one or other mutant residue without significant structural changes [Extended Data Figure 9]. Each mutation will add two negatively charged side chains per rung in the second shell around the central cavity, reducing the shell’s net positive charge. The presence of D51 or A53 may thus give rise to similar Type I and Type II filament structures, but the molecular composition of the density in the central cavity may be different from that in sporadic MSA. We cannot exclude that anionic, non-proteinaceous molecules in the central cavity are protective. It remains to be shown if they are cofactors, which can promote the assembly of α-synuclein into filaments.

By mass spectrometry of sarkosyl-insoluble α-synuclein from putamen, N-terminal acetylation, C-terminal truncation and ubiquitination of K6 and K12 were common to MSA cases 1-5. In the sequences of the filament cores, K21 was ubiquitinated in all cases. Despite identical Type I and Type II filament structures, sarkosyl-insoluble α-synuclein from only some cases of MSA also showed ubiquitination of K23, K60 and K80, acetylation of K21, K23, K32, K34, K45, K58, K60, K80 and K96, as well as phosphorylation of T39, T59, T64, T72 and T81. The percentage of α-synuclein molecules modified at a given residue is not known.

Ubiquitination of K80 was detected in sarkosyl-insoluble α-synuclein from MSA cases 2 and 5 (disease durations of 18 and 19 years), with a preponderance of Type II filaments. This bulky post-translational modification, which is compatible with the PF-IIA structure, clashes with the surroundings of the K80 side chain in PF-IA, PF-IB and PF-IIB. Moreover, one end of the peptide-like density, which is specific for Type II filaments, is located next to K80 of PF-IIA. Phosphorylation of T72 may favour PF-IIA over PF-IA. Whereas the side chain of T72 is buried in PF-IA, it borders a large cavity between the outer and central layers of PF-IIA. Phosphorylation of T81 may distinguish between PF-IIB_1_ and PF-IIB_2_, since the side chain of this residue is buried in PF-IIB_1_, but solvent-exposed in PF-IIB_2_. A post-translational modification in only one protofilament may favour the formation of asymmetrical Type I and Type II filaments. Thus, in the structures of PF-A, the side chain of K60 is solvent-exposed and can carry a bulky modification. In PF-B structures, in contrast, it is buried in the interfaces between protofilaments.

## DIFFERENT FILAMENT STRUCTURES MAY DEFINE SYNUCLEINOPATHIES

Our results show that α-synuclein filaments adopt the same structures in different individuals with MSA. Previously, we made similar observations for tau filaments from the brains of individuals with Alzheimer’s disease (4, 6), Pick’s disease (5), CTE (7) and CBD (9). Tau filaments adopt an identical fold in individuals with the same disease, but different tauopathies are characterised by distinct folds. To assess if the same is true of synucleinopathies, we used cryo-EM to examine α-synuclein filaments that were isolated from the brains of 3 individuals with a neuropathologically confirmed diagnosis of DLB.

In frontal cortex and amygdala, abundant Lewy bodies and Lewy neurites were stained by an antibody specific for α-synuclein phosphorylated at S129 [Extended Data Figure 10a]. Following sarkosyl extraction, α-synuclein filaments from the brains of individuals with DLB did not appear to twist and were thinner than those from the brains of individuals with MSA [Extended Data Figure 10b; Extended Data Figure 11a]. Similar differences between α-synuclein filaments from the brains of individuals with Lewy pathology and MSA have been described (23, 53). Moreover, unlike MSA, most sarkosyl-insoluble α-synuclein phosphorylated at S129 from DLB brains was SDS-insoluble, in agreement with previous findings (57). Immunogold negative-stain EM with anti-α-synuclein antibody PER4 showed decoration of DLB filaments [Extended Data Figure 10c], in confirmation of previous findings (59). The lack of twist precluded three-dimensional structure determination of α-synuclein filaments from DLB, but based on reference-free 2D class averages [Extended Data Figure 11b], we conclude that the structures of α-synuclein filaments of DLB are different from those of MSA.

Differences also exist with the tau filament structures determined thus far. Unlike MSA, each tauopathy is characterised by a single protofilament fold, with tau filaments consisting of one or two protofilaments (4-7,9). Distinct types of filaments are formed through variable inter-protofilament packing. With the exception of the straight filaments of Alzheimer’s disease, pairs of identical tau protofilaments are symmetrical.

This contrasts with MSA, where Type I and Type II filaments are formed by the asymmetrical packing of two pairs of different protofilaments, resulting in four distinct α-synuclein folds [Figure 2; Extended Data Figure 6]. Moreover, whereas the protofilament interfaces in tau filaments span at most 6 residues and are always between two identical monomers, the protofilament interfaces of α-synuclein filaments span more than 25 residues from each non-identical protofilament, and the N-terminal arms reach out to molecules in subsequent β-sheet rungs. This is bound to have an impact on seeded aggregation, as it results in differences in enthalpic and entropic gains, when a new molecule is incorporated into the filament.

## MSA FILAMENTS DIFFER FROM THOSE ASSEMBLED WITH RECOMBINANT ALPHA-SYNUCLEIN

We next compared the structures of α-synuclein filaments from MSA brains with those assembled *in vitro* from recombinant wild-type and mutant α-synucleins (60–67) [Figure 3d-f; Extended Data Figure 12].

The greatest differences are in the extended sizes of the MSA protofilaments and in their asymmetrical packing. None of the recombinant α-synuclein filaments contain the long N-terminal arms of MSA filaments and most recombinant filaments consist of either one protofilament or two identical protofilaments related by helical symmetry.

Like all MSA protofilaments, some recombinant α-synuclein protofilaments also contain three-layered L-shaped motifs [Figure 3d]. The first structure of recombinant α-synuclein filaments, 2N0A (60), belongs to this group. It described a Greek-key motif for residues G68-V95 of α-synuclein, which covers most of the NAC region. The conformation of the Greek-key motif is similar to that of the inner and central layers of the bodies of MSA filaments. The outer layer is also present in 2N0A, but is more disordered than in MSA protofilaments. One feature that recombinant α-synuclein filaments with the three-layered L-shaped motif have in common is that they were assembled in the presence of polyanions, like phosphate, or of chaotropic, negatively charged ions, such as bromine (60–63). The structural similarities between MSA and recombinant α-synuclein protofilaments are greatest in the residues that make up the outer and central layers, and in the turns between them.

The turn made by residues K43-V52 at the start of the outer layer of the L-shaped motif is nearly identical for protofilaments of MSA and those made using recombinant α-synuclein [Figure 3e]. In addition, the packing of the central layer against the start of the outer layer in multiple recombinant α-synuclein structures, including the formation of a salt bridge between E46 and K80, is similar to that in the structures of MSA protofilaments. It has been suggested that the additional densities in front of the side chains of K43, K45 from one protofilament, and of K58 from the other, correspond to phosphate ions (61, 64). Recombinant α-synuclein exhibits an increased propensity to aggregate upon exposure to anionic phospholipids (68, 69). Moreover, poly(adenosine 5’-diphosphate-ribose) has been shown to promote the formation of toxic strains of recombinant α-synuclein filaments (70).

The finding that in MSA Type I and Type II filaments the side chains of K43, K45 and H50 of both protofilaments point towards the strong additional density in the inter-protofilament cavity raises the possibility that this density may also correspond to molecules that contain phosphate groups. Unlike in the recombinant α-synuclein structures, the pseudo-symmetric cavity in MSA Type II filaments can accommodate approximately two phosphate groups per β-sheet rung, consistent with the size of the density [Figure 2; Extended Data Figure 6]. In Type I filaments, the central cavity is more open on one side and can therefore accommodate a larger density.

A second conformation that is nearly identical between MSA filaments and several recombinant α-synuclein structures is the turn between outer and central layers, which is formed by residues V63-T72 [Figure 3f]. Moreover, A69 and V71 on the outside of this turn pack against two residues on the outside of the inner layer through tight interdigitations of small side chains. In MSA PF-IA and PF-IIA filaments, as well as in 6OSM (60), these residues are A89 and A91; in MSA PF-IB and PF-IIB filaments, as well as in 6PEO, they are G93 and V95; in several recombinant α-synuclein structures, they are A91 and G93.

The bend in the central layer, the turn between central and inner layers, as well as the length of the inner layer, are more variable. Thus, despite topological similarities, in recombinant α-synuclein protofilaments, none of the three-layered L-shaped motifs are identical to those of MSA protofilaments [Extended Data Figure 12]. The closest similarity to an *in vitro* structure is between PF-IIB_2_ and 6PEO (65), which differ only in the bend positions in the outer layer (between E57 and K58 for PF-IIB_2_ and between T59 and K60 for 6PEO).

The finding that the structures of α-synuclein filaments from MSA differ from those of assembled recombinant proteins is consistent with the observation that rationally designed aggregation inhibitors of α-synuclein reduced aggregation by MSA and recombinant filament seeds to different extents (71). This finding is also reminiscent of similar results for tau filaments (4–9), even though marked differences exist between tau and α-synuclein. Recombinant tau is very soluble and requires cofactors to form filaments *in vitro*, whereas assembly of recombinant α-synuclein into filaments proceeds in the absence of cofactors. Moreover, α-synuclein exists as a single protein of 140 amino acids, whereas 6 tau isoforms with lengths ranging from 352-441 amino acids are expressed in adult human brain; the isoform composition of filaments varies among some tauopathies (72). Still, as was the case of recombinant α-synuclein filaments (60–67), the structures of heparin-induced filaments of recombinant tau (8) differed from those present in disease (4-7,9). In both cases, *in vitro* assembled filaments were smaller and adopted topologically simpler conformations.

## OUTLOOK

Our results establish the presence of two distinct, but similar, types of α-synuclein filaments in MSA, and suggest that different conformers (or strains) of assembled α-synuclein exist in DLB. To understand the causes and spreading of α-synuclein pathology, as well as what distinguishes synucleinopathies. it will be important to identify the mechanisms of seed formation and subsequent assembly.

Despite overall differences between α-synuclein filaments in MSA and those assembled from recombinant protein *in vitro*, the latter may provide useful insights. When compared to MSA filaments, they exhibit both overall structural diversity and topological similarity in the three-layered, L-shaped motif. We therefore hypothesise that α-synuclein molecules may exist in a cloud (73) of similar conformations, prior to their incorporation into filaments. The three-layered L-shaped topology could be driven by formation of the conserved turn between outer and central layers and its packing against the inner layer, as well as by the packing of the turn at the start of the inner layer against the end of the central layer. The latter may be stabilised further by the formation of a salt bridge between E46 and K80.

Even if α-synuclein monomers or oligomers adopt a cloud of different conformations early on, our structures show that only two types of filaments are present in end-stage MSA brains. The reasons for this compelling structural specificity, which was also observed in tauopathies (4-7,9), are unclear. The presence of post-translational modifications in assembled α-synuclein is well established, but their relevance for assembly is not understood (1). In addition, the structures of α-synuclein filaments from MSA reveal the presence of non-proteinaceous molecules, reminiscent of our findings in tauopathies (7, 9). It will be important to identify the chemical nature of these molecules and to study their effects, alone or in conjunction with post-translational modifications, on α-synuclein and tau assembly. Understanding the structural specificity of filament assembly in disease will facilitate the development of tracers for imaging filamentous amyloid assemblies of α-synuclein in brain, as well as of molecules that prevent, inhibit and reverse filament formation.

## Acknowledgements

We thank the patients’ families for donating brain tissues; T. Nakane for help with RELION; T. Darling and J. Grimmett for help with high-performance computing; F. Epperson, U. Kuederli, R. Otani and R.M. Richardson for support with neuropathology; R.A. Crowther, B. Falcon, S. Lovestam, M.G. Spillantini and W. Zhang for helpful discussions. M.G. is an Honorary Professor in the Department of Clinical Neurosciences of the University of Cambridge and an Associate Member of the U.K. Dementia Research Institute. This work was supported by the U.K. Medical Research Council (MC_UP_A025_1013, to S.H.W.S., and MC_U105184291, to M.G.), Eli Lilly and Company (to M.G.), the European Union (EU/EFPIA/Innovative Medicines Initiative [2] Joint Undertaking IMPRIND, project 116060, to M.G.), the Japan Agency for Medical Research and Development (JP18ek0109391 and JP18dm020719, to M.H.), the U.S. National Institutes of Health (P30-AG010133 and U01-NS110437, to B.G.) and the Department of Pathology and Laboratory Medicine, Indiana University School of Medicine (to B.G.). We acknowledge Y. Chaban at Diamond for access to and support from the cryo-EM facilities at the U.K. electron Bio-Imaging Centre (eBIC), proposal EM17434-75, funded by the Wellcome Trust, MRC and the BBSRC, for acquisition of the MSA case 1 dataset. This study was supported by the MRC-LMB EM facility.

## Author contributions

A.T., B.G., T.M., T.T., K.H., S.M., M.Y. and M.H. identified patients, performed neuropathology and extracted α-synuclein filaments from MSA cases 1-5 and DLB cases 1-2; M.S. extracted α-synuclein filaments from DLB case 3 and conducted immunolabelling of filaments from MSA cases 1-5 and DLB cases 1-3; F.K. and M.H. carried out mass spectrometry; A.T. and M.H. did seeded aggregation; M.S. and Y.S. performed cryo-EM; Y.S., M.S. and S.H.W.S. analysed the cryo-EM data; Y.S. and A.G.M. built the atomic models; S.H.W.S. and M.G. supervised the project; all authors contributed to writing the manuscript.

## METHODS

No statistical methods were used to predetermine sample size. The experiments were not randomized and investigators were not blinded to allocation during experiments and outcome assessment.

### Clinical history and neuropathology

MSA case 1 was an 85-year-old woman who died with a neuropathologically confirmed diagnosis of MSA-P following a 9-year history of bradykinesia, rigidity in upper and lower limbs and autonomic failure. MSA case 2 was a 68-year-old woman who died with a neuropathologically confirmed diagnosis of MSA-C following an 18-year history of cerebellar ataxia, gait disturbance and autonomic failure. MSA case 3 was a 59-year-old man who died with a neuropathologically confirmed diagnosis of MSA-C following a 9-year history of dysarthria, cerebellar ataxia and autonomic failure. MSA case 4 was a 64-year-old man who died with a neuropathologically confirmed diagnosis of MSA-C following a 10-year history of cerebellar ataxia, dysarthria and autonomic failure. MSA case 5 was a 70-year-old man who died with a neuropathologically confirmed diagnosis of MSA-C following a 19-year history of cerebellar ataxia and autonomic failure. DLB case 1 was a 59-year old man who died with a neuropathologically confirmed diagnosis of DLB following a 10-year history of resting tremor, bradykinesia, rigidity, postural instability and visual hallucinations. DLB case 2 was a 74-year old man who died with a neuropathologically confirmed diagnosis of diffuse Lewy body disease following a 13-year history of bradykinesia, postural instability and visual hallucinations. DLB case 3 was a 78-year old man who died with a neuropathologically confirmed diagnosis of diffuse Lewy body disease following a 15-year history of resting tremor, bradykinesia, autonomic symptoms and visual hallucinations.

### Extraction of α-synuclein filaments

Sarkosyl-insoluble material was extracted from fresh-frozen brain regions of MSA and DLB, essentially as described (53). Briefly, tissues were homogenised in 20 vol. (v/w) extraction buffer consisting of 10 mM Tris-HCl, pH 7.5, 0.8 M NaCl, 10% sucrose and 1 mM EGTA. Homogenates were brought to 2% sarkosyl and incubated for 30 min. at 37° C. Following a 10 min. centrifugation at 10,000 g, the supernatants were spun at 100,000 g for 20 min. The pellets were resuspended in 500 μl/g extraction buffer and centrifuged at 3,000 g for 5 min. The supernatants were diluted 3-fold in 50 mM Tris-HCl, pH 7.5, containing 0.15 M NaCl, 10% sucrose and 0.2% sarkosyl, and spun at 166,000 g for 30 min. Sarkosyl-insoluble pellets were resuspended in 100 μl/g of 30 mM Tris-HCl, pH 7.4. We used approximately 0.5 g tissue for cryo-EM and 0.5 g for negative stain immuno-EM In some experiments, sarkosyl-insoluble pellets were resuspended in 30 mM Tris-HCl, 2% SDS, left at room temperature for 30 min. and spun at 100,000 g for 30 min The pellets were resuspended in 8M urea. Both supernatants and pellets were immunoblotted using anti-pS129 α-synuclein antibody.

### Immunolabelling and histology

Immunogold negative-stain EM and Western blotting were carried out as described (74). Using the same procedure as above, filaments were extracted from putamen of MSA cases 1-5, cerebellum of MSA case 1, frontal cortex of MSA case 3, frontal cortex of DLB cases 1 and 2, and amygdala of DLB case 3. PER4 (59) was used at 1:50 (a polyclonal rabbit serum that recognises the carboxy-terminal region of α-synuclein). Images were acquired at x11,000 with a Gatan Orius SC200B CCD detector on a Tecnai G2 Spirit at 120 eV. For Western blotting, the samples were resolved on 4-12% Bis-Tris gels (NuPage) and the primary antibodies diluted in PBS plus 0.1% Tween 20 and 5% non-fat dry milk. Prior to blocking, membranes were fixed with 1% paraformaldehyde for 30 min. Primary antibodies were: Syn303 (BioLegend) (75) at 1:4,000 (a monoclonal mouse antibody that recognises the amino-terminus of α-synuclein), PER4 at 1:4,000 and pS129 (ab51253, Abcam) at 1:5,000 (a monoclonal rabbit antibody that recognises α-synuclein phosphorylated at S129).

Histology and immunohistochemistry were carried out as described (53, 76). Some brain sections (8 μm) were counterstained with haematoxylin. The primary antibody was specific for α-synuclein phosphorylated at S129 (ab51253).

### Seeded α-synuclein aggregation

The ability of sarkosyl-insoluble fractions from the putamen of MSA cases 1-5 to convert expressed soluble α-synuclein into its abnormal form was examined, as described (77, 78). Following addition of variable amounts of seeds (ranging from 1-4,700 pg/ml), transfected cells were incubated for 3 days. Sarkosyl-insoluble α-synuclein was extracted, run on 15% SDS-PAGE and immunoblotted with a mouse monoclonal antibody specific for α-synuclein phosphorylated at S129 (pSyn64 at 1:1,000) (58). Band intensities were quantified using ImageJ software.

### Mass spectrometry of sarkosyl-insoluble α-synuclein

Protease digestion and nano-flow liquid chromatography-ion trap mass spectrometry (LC-MS/MS, Thomas Scientific, Q Exactive HF) were used to identify post-translational modifications in sarkosyl-insoluble α-synuclein extracted from the putamen of MSA cases 1-5 (79). The concentration of α-synuclein was determined using an ELISA kit (Abcam). Sarkosyl-insoluble fractions containing approximately 65 ng of α-synuclein were treated with 70% formic acid for 1 h at room temperature, diluted in water and dried. They were digested overnight with trypsin/lysyl-endopeptidase. Peptides were then analysed by LC-MS/MS.

### Electron cryo-microscopy

Extracted tau filaments were applied to glow-discharged holey carbon gold grids (Quantifoil R1.2/1.3, 300 mesh) and plunge-frozen in liquid ethane using an FEI Vitrobot Mark IV. Micrographs were acquired on a Gatan K2 Summit direct detector in counting mode or a Gatan K3 direct detector in super-resolution mode on a Thermo Fischer Titan Krios microscope that was operated at 300 kV. Inelastically scattered electrons were removed by a GIF Quantum energy filter (Gatan) using a slit width of 20 eV. Further details are given in Extended Data Table 1.

### Helical reconstruction

Movie frames were corrected for beam-induced motion and dose-weighted using RELION’s motion-correction implementation (80). Aligned, non-dose-weighted micrographs were then used to estimate the contrast transfer function using CTFFIND-4.1 (81). All subsequent image-processing steps were performed using helical reconstruction methods in RELION 3.0 (82). Filaments were picked manually.

### MSA datasets

Segments for reference-free 2D classification comprising an entire helical crossover were extracted using an inter-box distance of 14.1 Å. For samples extracted from putamen, segments with a box size of 750 pixels and a pixel size of 1.15 Å were downscaled to 256 pixels for MSA cases 2-5, and segments with a box size of 900 pixels and a pixel size of 0.83 Å were downscaled to 300 pixels for MSA case 1. For samples extracted from cerebellum of MSA case 1 and frontal cortex of MSA case 3, segments with a box size of 750 pixels and a pixel size of 1.15 Å were downscaled to 256 pixels. MSA Type I and Type II filaments from putamen were initially separated by reference-free 2D classification and segments contributing to suboptimal 2D class averages were discarded. For both types of filaments, an initial helical twist of −1.4° was calculated from the apparent crossover distances of filaments in micrographs, and the helical rise was fixed at 4.7 Å. Using these values, initial 3D models for both types were constructed *de novo* from 2D class averages of segments that comprise entire helical crossovers using the *relion_helix_inimodel2d* program (83). Type I and Type II filament segments were then re-extracted using box sizes of 220 pixels for MSA cases 2-5 or 320 pixels for MSA case 1, without downscaling. Starting with these segments and an initial *de novo* model low-pass filtered to 10 Å, 3D auto-refinement was carried out for several rounds with optimisation of helical twist and rise after reconstructions showed separation of β-strands along the helical axis. We then performed Bayesian polishing and contrast transfer function refinement, followed by 3D classification with local optimisation of helical twist and rise, but without further image alignment, to remove segments that yielded suboptimal 3D reconstructions. To further separate the subtypes of Type II filaments, segments of Type II filaments from MSA case 2 were subjected to additional supervised and focused 3D classifications of K45-V95 from PF-IIB; MSA Type II_1_ and Type II_2_ filaments served as references. For all cases, selected segments were used for further 3D auto-refinement. Final reconstructions were sharpened using the standard post-processing procedures in RELION (82). Overall resolution estimates were calculated from Fourier shell correlations at 0.143 between the two independently refined half-maps, using phase-randomisation to correct for convolution effects of a generous, soft-edged solvent mask that extended to 20 % of the height of the box. Local resolution estimates were obtained using the same phase-randomisation procedure, but with a soft spherical mask that was moved over the entire map. Using the *relion_helix_toolbox* program (83), helical symmetries were imposed on the post-processed maps, which were then used for model building. The reported ratios of MSA Type I and Type II filament segments in each case were determined by 2D classification of mixed sets of segments, which were re-extracted with box sizes of 750 or 900 pixels, while keeping the alignment parameters fixed to those resulting from the initial 3D refinements.

### DLB datasets

DLB filament segments were extracted using an inter-box distance of 14.1 Å. For DLB cases 1-3, segments with a box size of 540 pixels and a pixel size of 1.15 Å were downscaled to 270 pixels. Reference-free 2D classification was performed using standard procedures.

### Model building and refinement

Atomic models for Type I and Type II filaments were built *de novo* in Coot (84), using the maps of MSA case 1 and MSA case 2, respectively. Model building was started from the topologically conserved C-terminal bodies using the cryo-EM structure of recombinant α-synuclein filaments (6A6B) as an initial reference (60). For turns with weaker densities, models were built at low display thresholds.Models containing 5 β-sheet rungs were refined in real-space by PHENIX using local symmetry and geometry restraints (85). MolProbity (86) was used for model validation. Additional details are given in Extended Data Table 1.

### Ethical review processeses and informed consent

The studies carried out at Tokyo Metropolitan Institute of Medical Science and at Indiana University were approved through the ethical review processes of each Institution. Informed consent was obtained from the patients’ next of kin.

### Data availability

Cryo-EM maps have been deposited in the Electron Microscopy Data Bank (EMDB) under accession numbers EMD-10650 for Type I filaments from MSA case 1, EMD-10651 for Type II_1_ filaments from MSA case 2 and EMD-10652 for Type II_2_ filaments from MSA case 2. The corresponding atomic models have been deposited in the Protein Data Bank (PDB) under the following accession numbers: 6XYO for Type I filaments from MSA case 1, 6XYP for Type II_1_ filaments from MSA case 2 and 6XYQ for Type II_2_ filaments from MSA case 2.

## EXTENDED DATA FIGURE LEGENDS

**Extended Data Figure 1.**
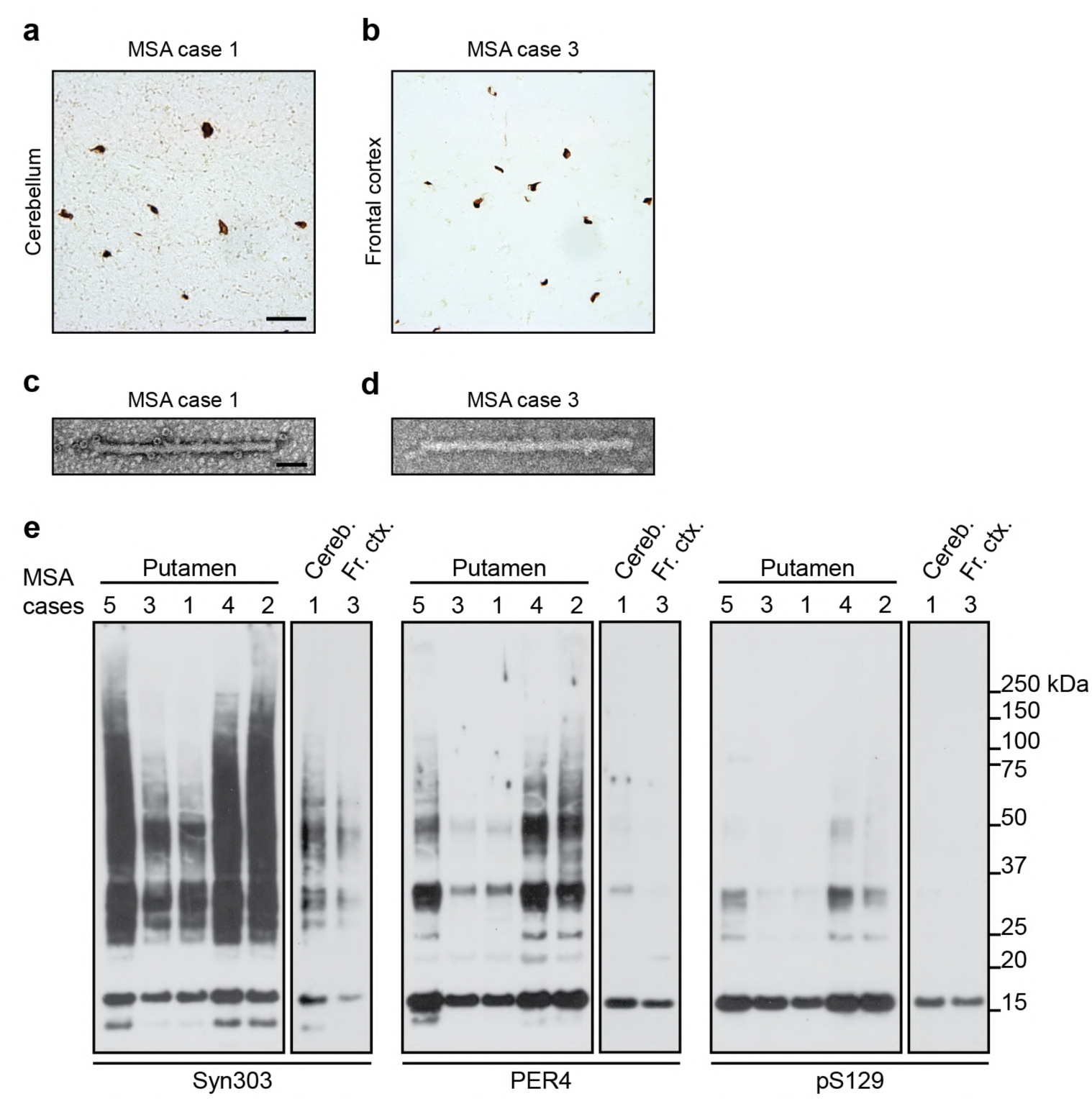
Filamentous α-synuclein pathology in MSA. (a,b) Staining of inclusions in cerebellum of MSA case 1 and frontal cortex of MSA case 3 by an antibody specific for α-synuclein phosphorylated at S129 (brown). Scale bar, 50 μm. (c,d), Negative-stain EM of filaments from cerebellum of MSA case 1 and frontal cortex of MSA case 3. Scale bar, 50 nm. (e), Immunoblots of sarkosyl-insoluble material from putamen of MSA cases 1-5, cerebellum of MSA case 1 and frontal cortex of MSA case 3, using anti-α-synuclein antibodies Syn303 (N-terminus), PER4 (C-terminus) and pS129 (phosphorylation of S129).

**Extended Data Figure 2.**
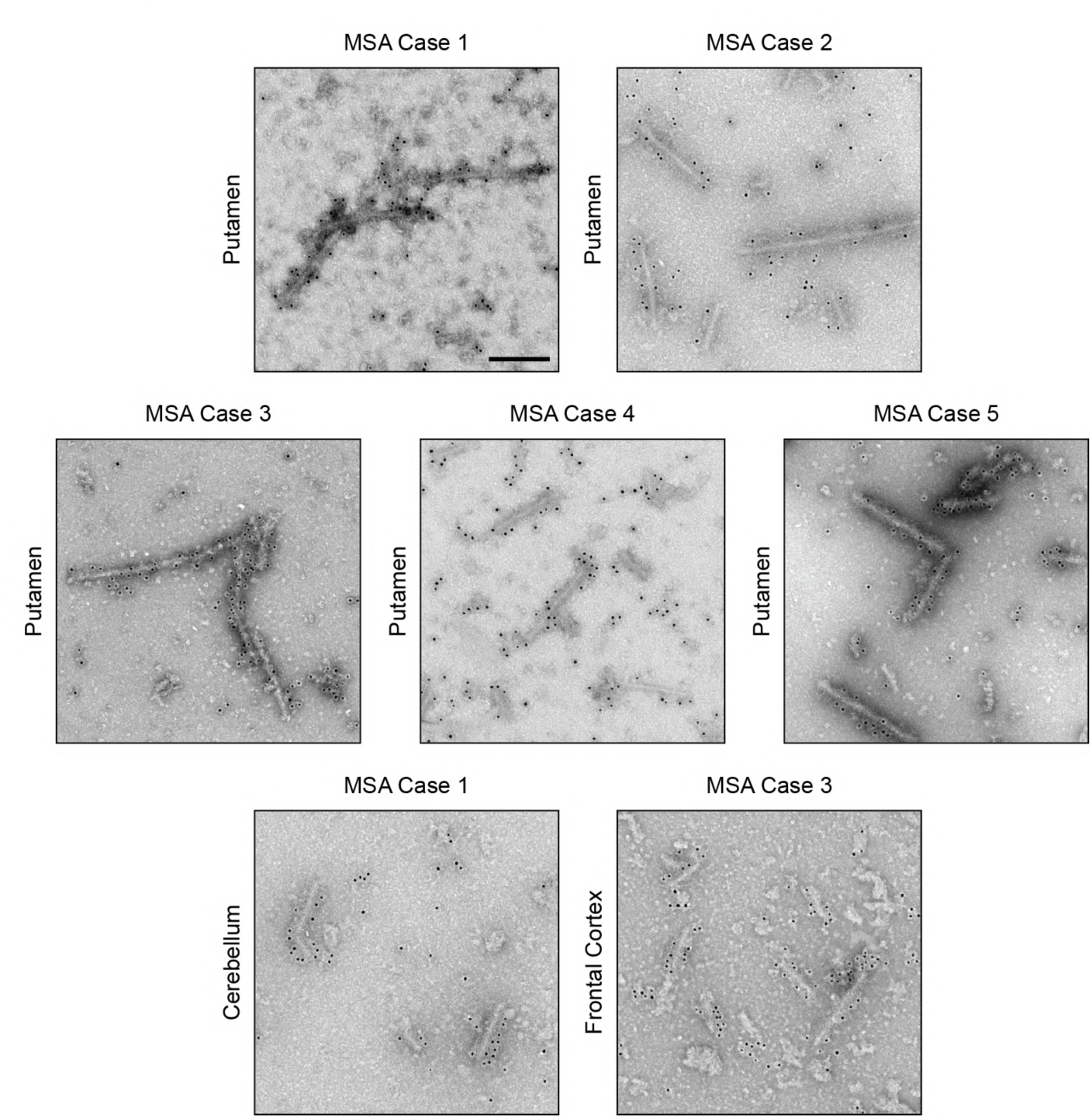
Immunolabelling of α-synuclein filaments extracted from MSA brains. Representative immunogold negative-stain EM images of α-synuclein filaments extracted from putamen of MSA cases 1-5, cerebellum of MSA case 1 and frontal cortex of MSA case 3. Filaments were labelled with antibody PER4, which recognises the C-terminus of α-synuclein. Scale bar, 200 nm.

**Extended Data Figure 3.**
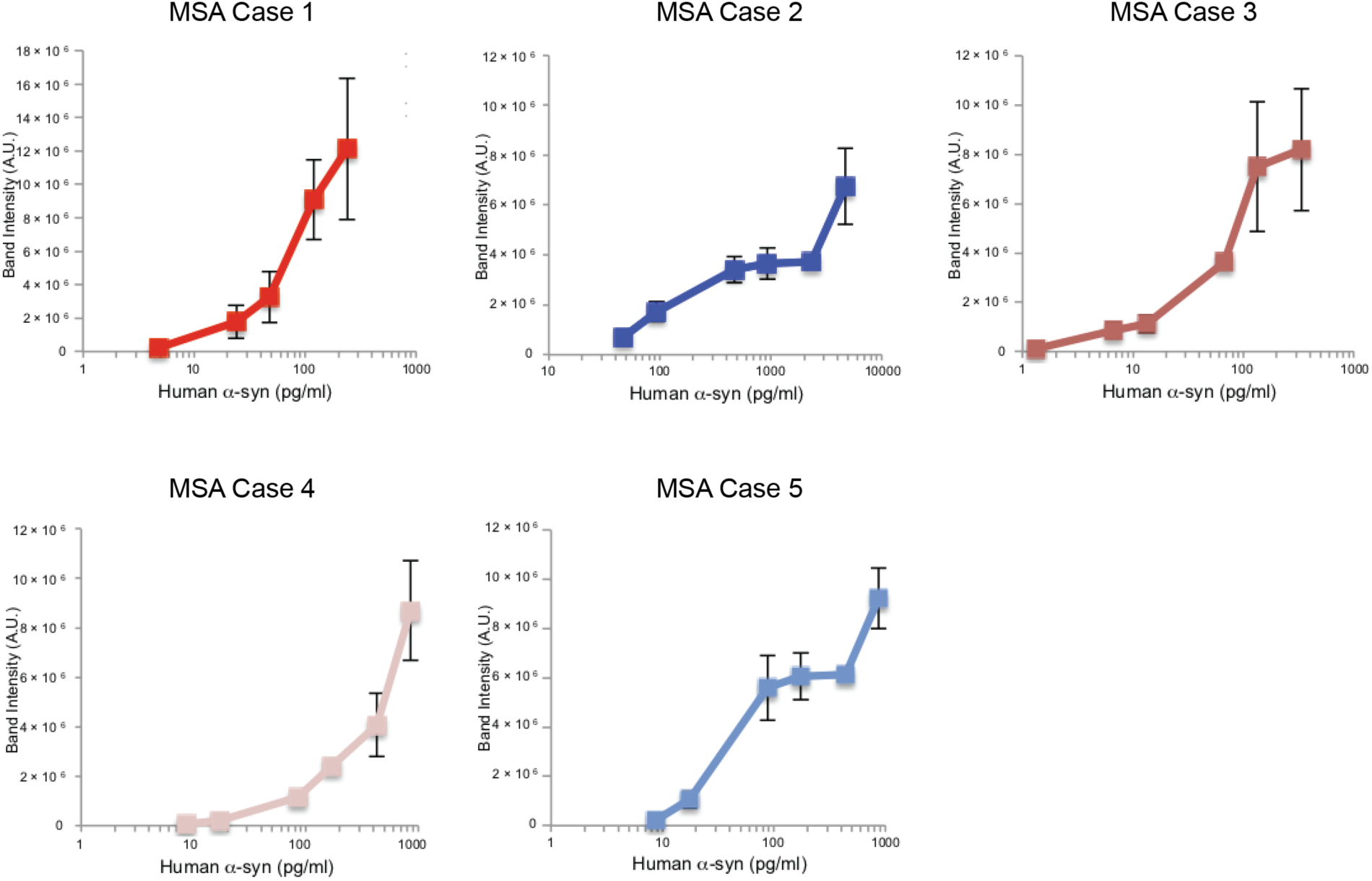
Aggregation of wild-type human α-synuclein in SH-SY5Y cells following addition of seeds from MSA cases 1-5. Quantitation of α-synuclein phosphorylated at S129 in SH-SY5Y cells following addition of variable amounts of α-synuclein seeds from the putamen of MSA cases 1-5. The results are expressed as means ± S.E.M. (n=3).

**Extended Data Figure 4.**
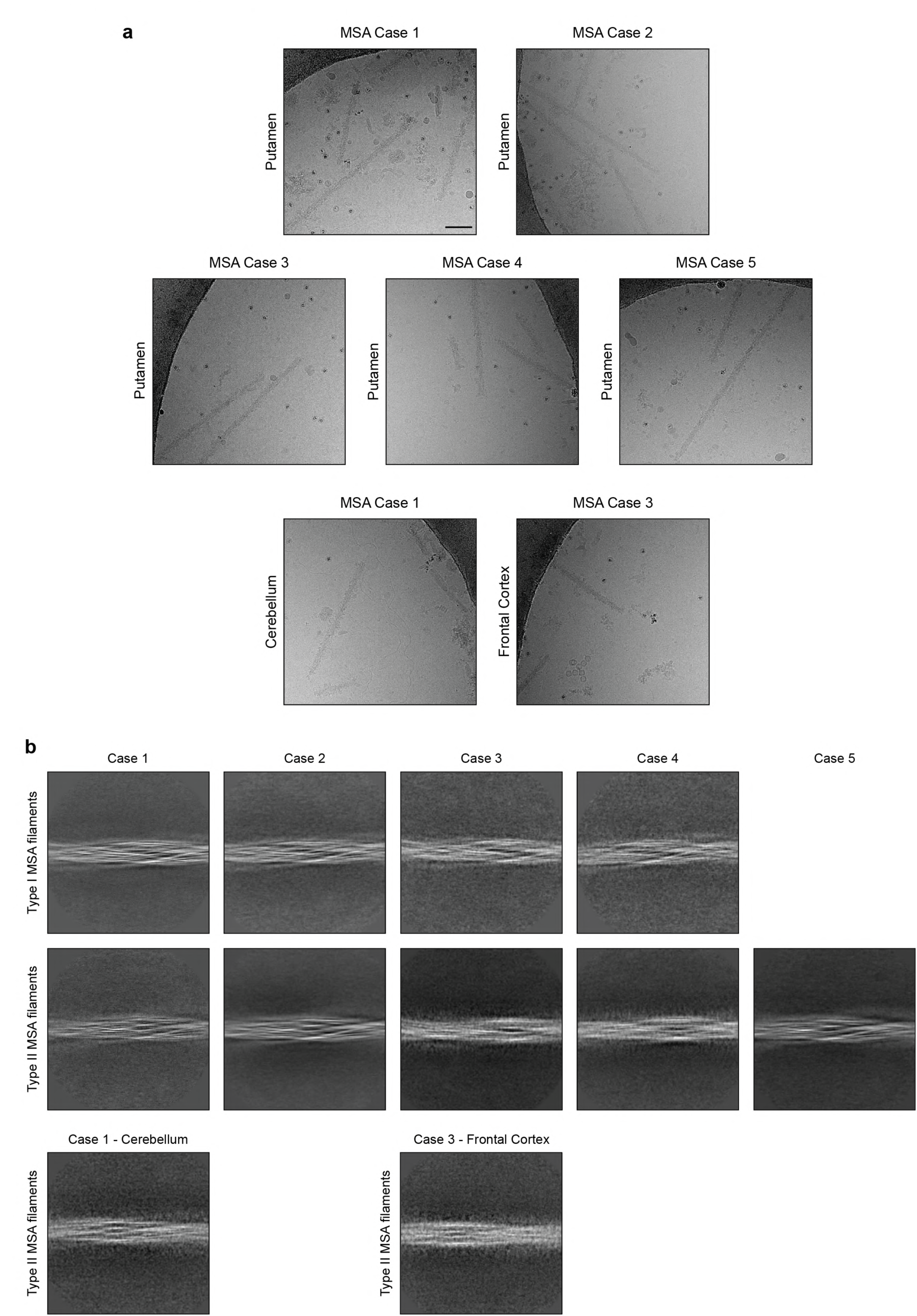
Cryo-EM images and 2D classification of MSA filaments. (a), Representative cryo-EM images of α-synuclein filaments from the putamen of MSA cases 1-5, cerebellum of MSA case 1 and frontal cortex of MSA case 3. Scale bar, 100 nm. (b), Two-dimensional class averages spanning an entire crossover of Type I and Type II α-synuclein filaments extracted from putamen of MSA cases 1-5, cerebellum of MSA case 1 and frontal cortex of MSA case 3.

**Extended Data Figure 5.**
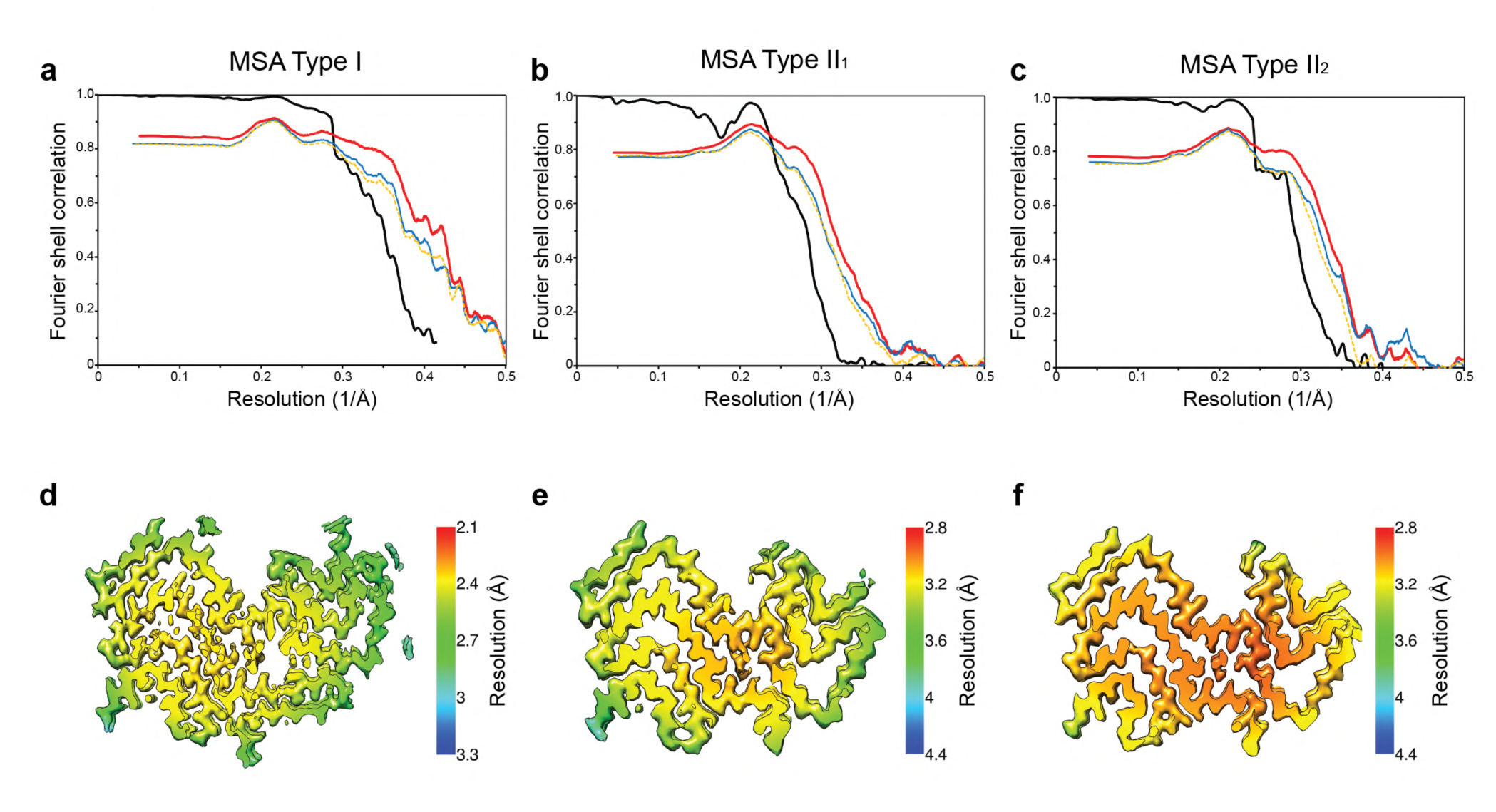
Resolution evaluation of cryo-EM maps and of reconstructed models. (a-c), For MSA Type I (a), Type II_1_ (b) and Type II_2_ (c) filaments: Fourier shell correlation (FSC) curves of two independently refined half-maps (black line); FSC curves of final cryo-EM reconstruction and refined atomic model (red); FSC curves of first half-map and the atomic model refined against this map (blue); FSC curves of second half-map and the atomic model refined against the first half-map (yellow dashes). (d-f), Local resolution estimates of the reconstructions of MSA Type I (d), Type II_1_ (e) and Type II_2_ (f) filaments.

**Extended Data Figure 6.**
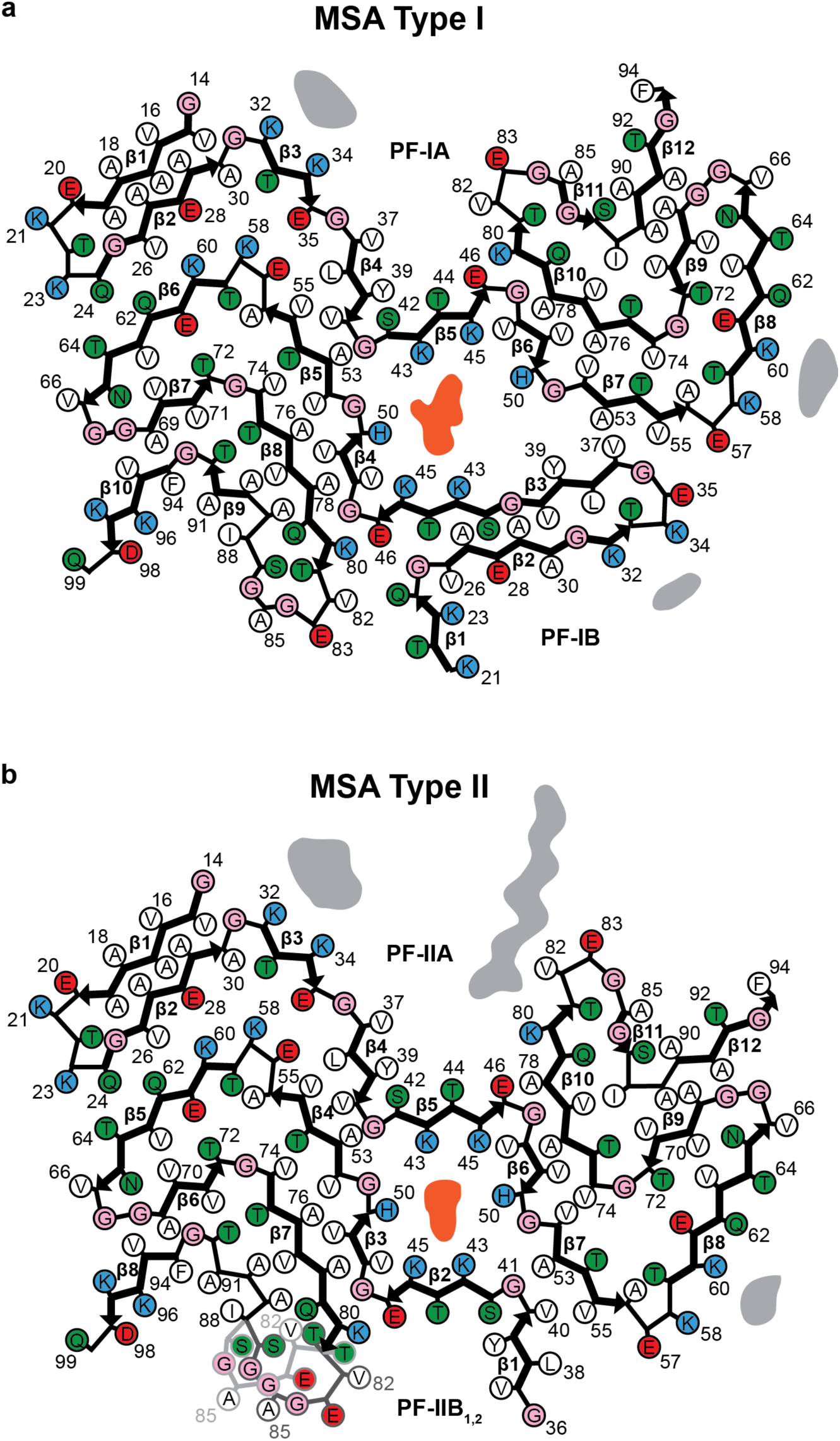
MSA Type I and Type II α-synuclein filaments. (a), Schematic of MSA Type I α-synuclein filament, showing asymmetric protofilaments IA and IB. The non-proteinaceous density at the protoflament interface is shown in light red. (b), Schematic of MSA Type II α-synuclein filament, showing asymmetric protofilaments IIA and IIB. The non-proteinaceous density at the protofilament interface is shown in light red.

**Extended Data Figure 7.**
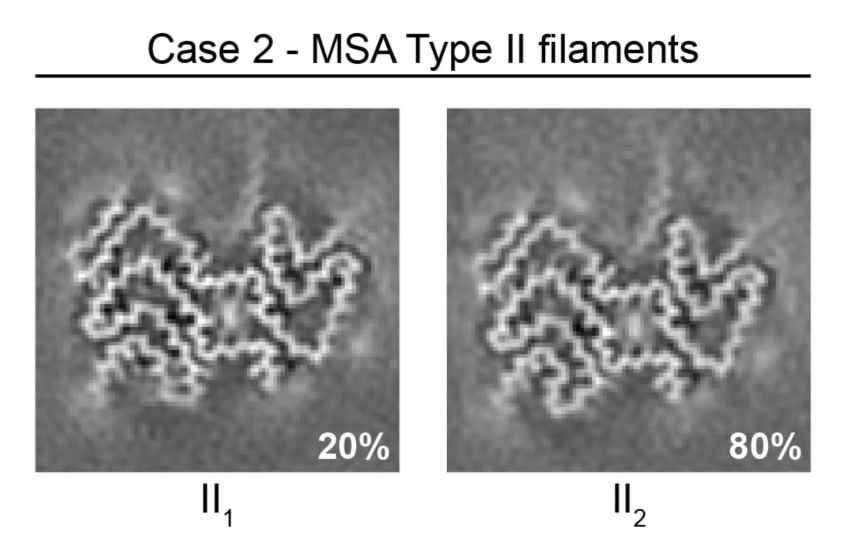
Cryo-EM maps of MSA Type II_1_ and Type II_2_ filaments. Type II filaments were extracted from the putamen of MSA case 2.

**Extended Data Figure 8.**
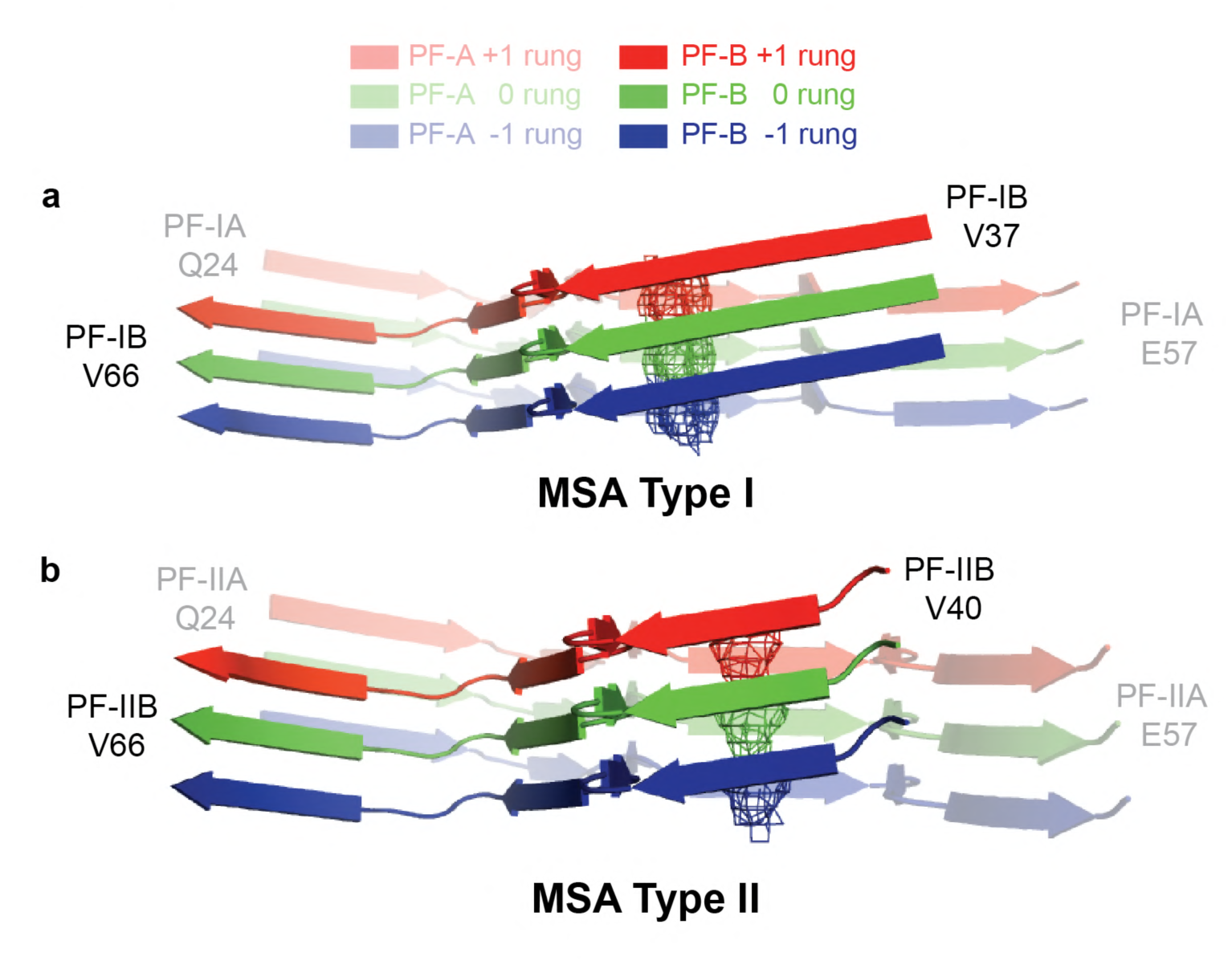
Differences in height of inter-protofilament interfaces of MSA Type I and Type II α-synuclein filaments. Rendered view of secondary structure elements in MSA Type I (a) and Type II (b) protofilament folds perpendicular to the helical axis of the inter-protofilament interfaces, depicted as three rungs.

**Extended Data Figure 9.**
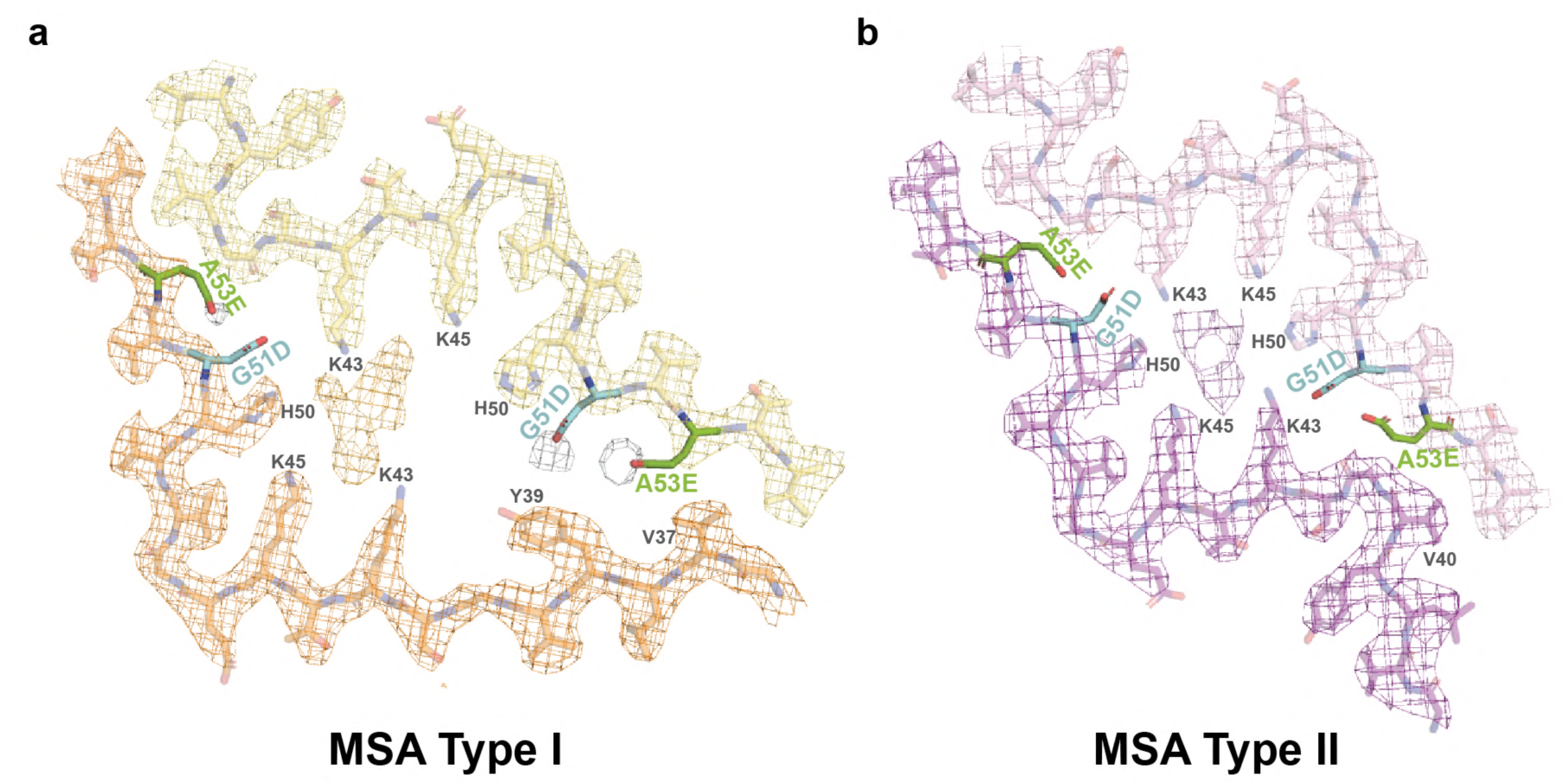
Compatibility of mutant α-synuclein (G51D or A53E) with MSA Type I and Type II filament folds. (a,b), Close-up views of atomic models of MSA Type I (a) and MSA Type II (b) α-synuclein folds containing D51 (cyan) and E53 (green).

**Extended Data Figure 10.**
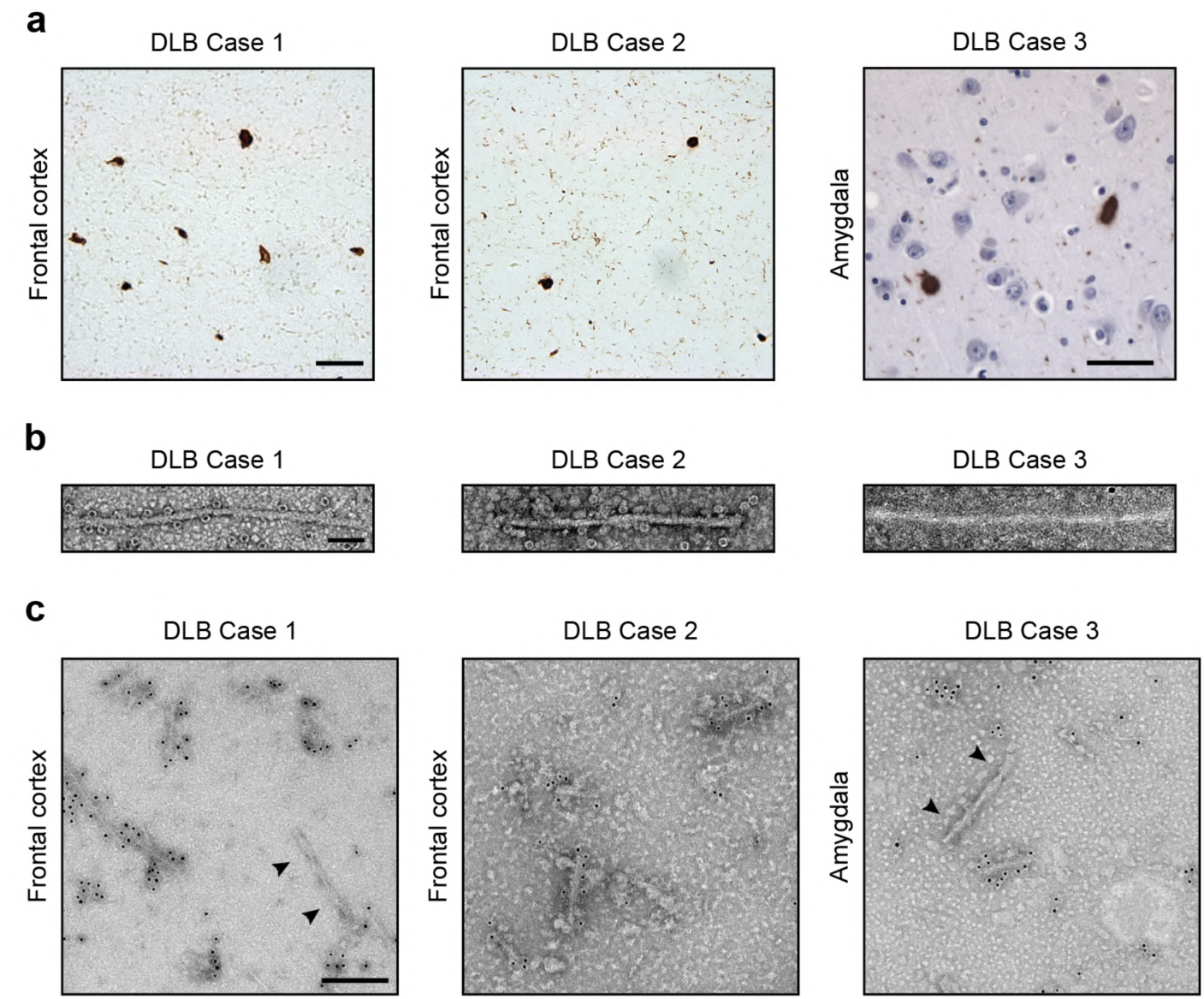
Filamentous α-synuclein pathology in DLB. (a), Staining of inclusions in frontal cortex of DLB cases 1 and 2 and amygdala of DLB case 3 by an antibody specific for α-synuclein phosphorylated at S129 (brown). Scale bar, 50 μm. (b), Negative-stain electron micrographs of filaments from frontal cortex of DLB cases 1 and 2 and amygdala of DLB case 3. Scale bar, 50 nm. (c), Representative immunogold negative-stain EM images of α-synuclein filaments extracted from frontal cortex of DLB cases 1 and 2 and amygdala of DLB case 3. Filaments were labelled with antibody PER4, which recognises the C-terminus of α-synuclein. Arrowheads point to an unlabelled tau paired helical filament. Scale bar, 200 nm.

**Extended Figure 11.**
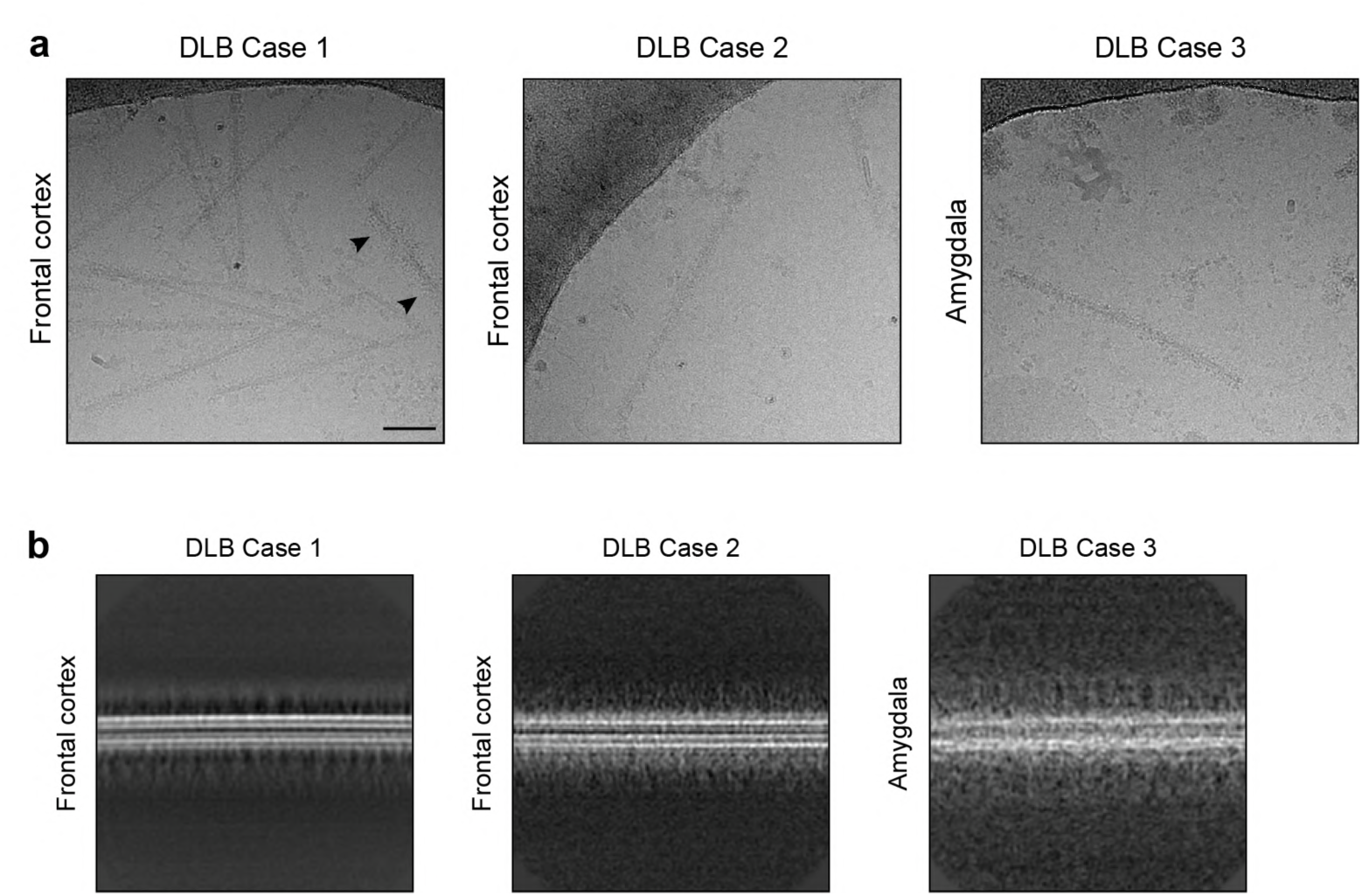
Cryo-EM images and 2D classification of DLB filaments. (a), Representative cryo-EM images of α-synuclein filaments from frontal cortex of DLB cases 1 and 2, and amygdala of DLB case 3. Arrowheads point to a tau paired helical filament. (b), Two-dimensional class averages of α-synuclein filaments extracted from frontal cortex of DLB cases 1 and 2 and amygdala of DLB case 3.

**Extended Data Figure 12.**
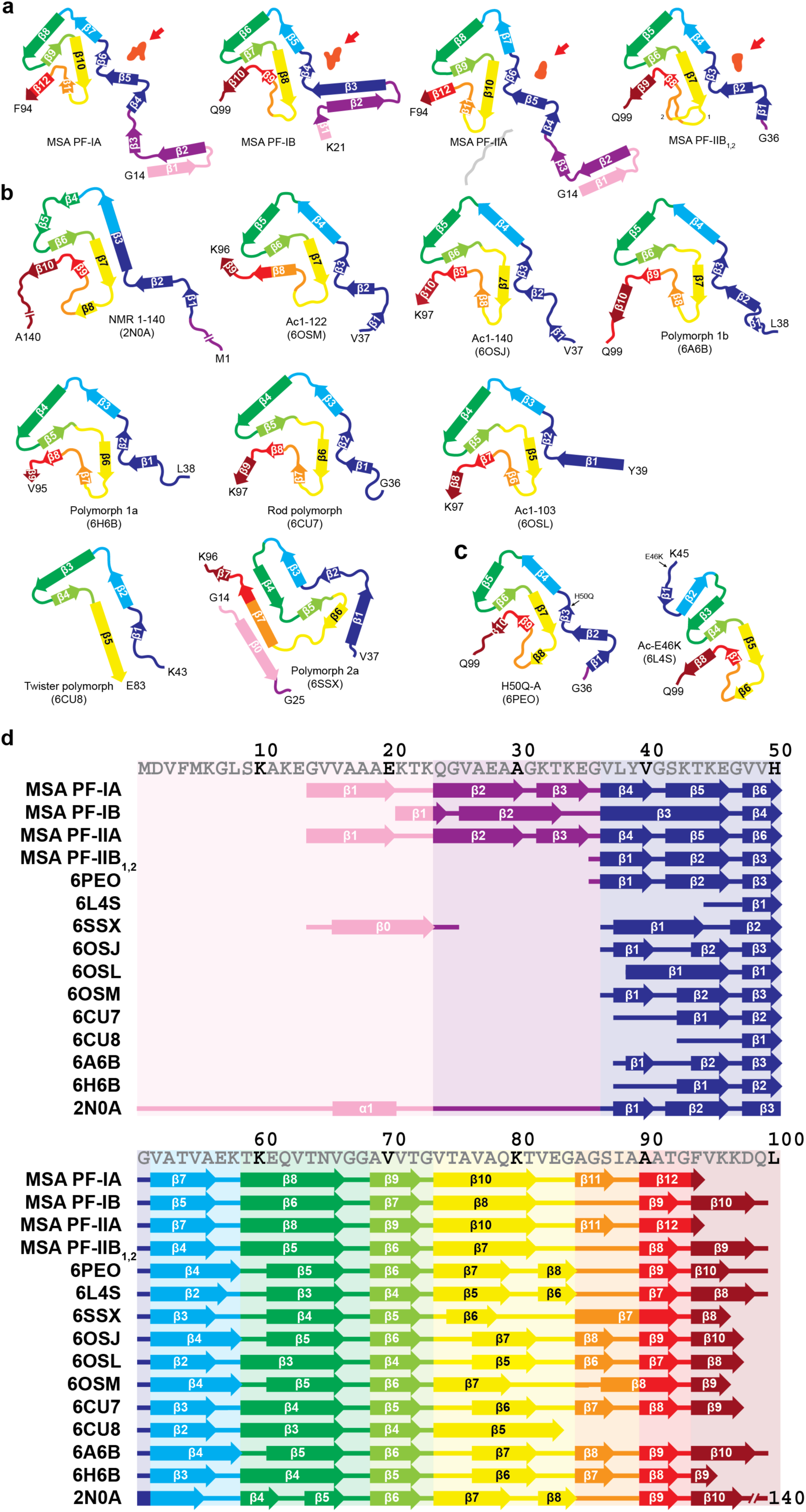
Structures of α-synuclein protofilament cores. (a), Schematic of secondary structure elements in the α-synuclein protofilament cores of MSA. Red arrows point to the non-proteinaceous density (in light red) at protofilament interfaces. (b,c), Secondary structure elements in the α-synuclein protofilament cores assembled from recombinant wild-type (b) and mutant (c) α-synuclein *in vitro*. (d), Schematic depicting the first 100 amino acids of human α-synuclein, with the β-strands found in the cores of α-synuclein protofilaments (PF) marked by thick arrows.

**Extended Data Table 1.**
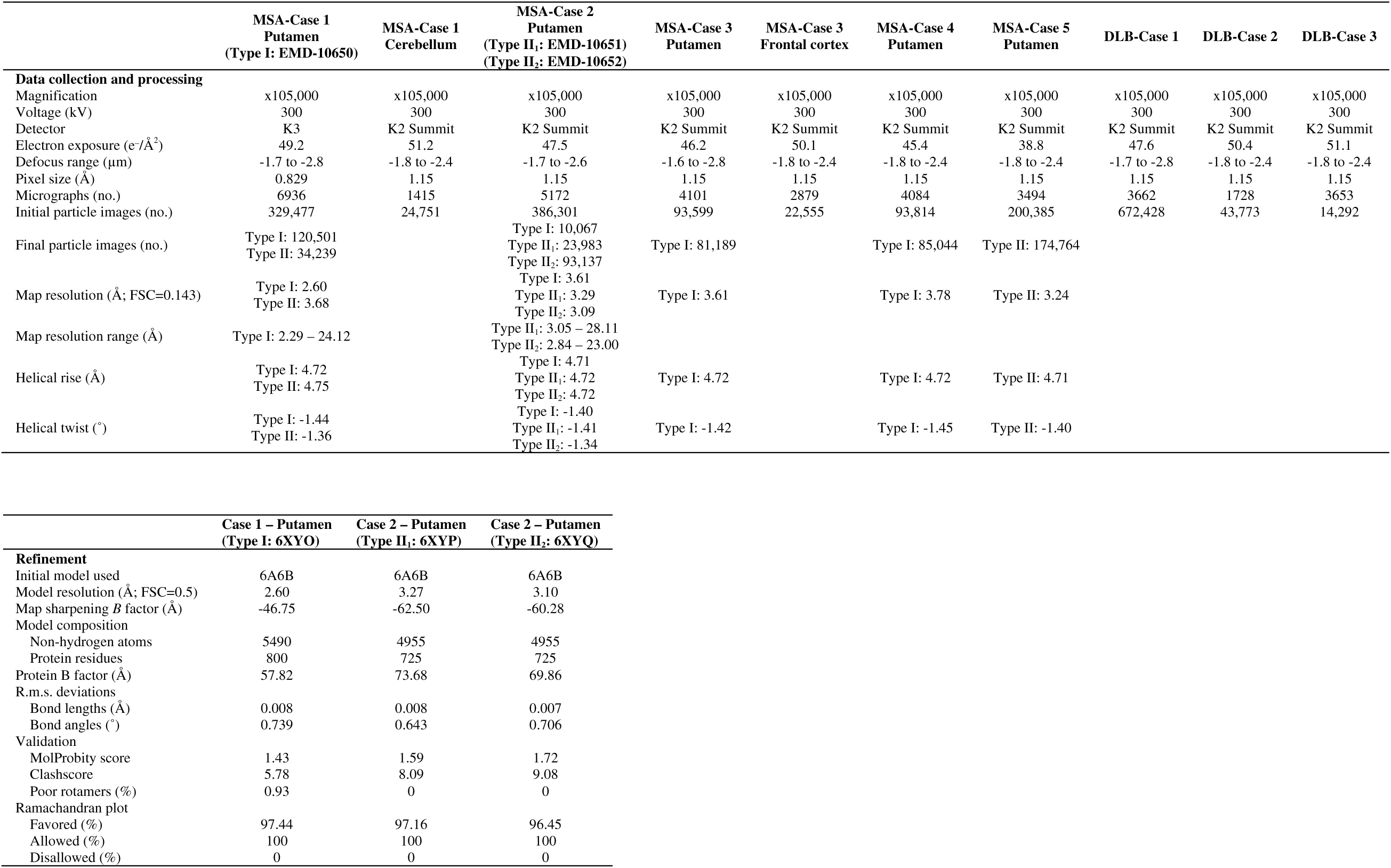
Cryo-EM data collection, refinement and validation statistics.

